# The hydroalcoholic extract of *Uncaria tomentosa* (Cat’s claw) inhibits the replication of novel coronavirus (SARS-CoV-2) *in vitro*

**DOI:** 10.1101/2020.11.09.372201

**Authors:** Andres F. Yepes-Perez, Oscar Herrera-Calderón, Lizdany Flórez-Álvarez, María I. Zapata-Cardona, Lina Yepes, Wbeimar Aguilar, María T. Rugeles, Wildeman Zapata

**Author notes:** **Corresponding author.** Jr. Puno 1002, Faculty of Pharmacy and Biochemistry, Universidad Nacional Mayor de San Marcos, Lima 15001, Peru.

## Abstract

The coronavirus disease 2019 (COVID-19) has become a serious problem for public health since it was identified in the province of Wuhan (China) and spread around the world producing high mortality rates and economic losses. Nowadays, WHO recognizes traditional, complementary, and alternative medicine for treating COVID-19 symptoms. Therefore, we investigated the antiviral potential of the hydroalcoholic extract of *Uncaria tomentosa* stem bark from Peru against SARS-CoV-2 *in vitro*. The antiviral activity of *U. tomentosa* against SARS-CoV-2 *in vitro* was assessed in Vero E6 cells using cytopathic effect (CPE) and plaque reduction assay. After 48h of treatment, *U. tomentosa* showed an inhibition of 92.7 % of SARS-CoV-2 at 25.0 µg/mL (p<0.0001) by plaque reduction assay on Vero E6 cells. In addition, *U. tomentosa*, induced a reduction of 98.6 % (p=0.02) and 92.7 % (p=0.03) in the CPE caused by SARS-CoV-2 on Vero E6 cells at 25 µg/mL and 12.5 µg/mL, respectively. The EC_50_ calculated for *U. tomentosa* extract by plaque reduction assay was 6.6 µg/mL (4.89 – 8.85 µg/mL) for a selectivity index of 4.1. The EC_50_ calculated for *U. tomentosa* extract by TCID_50_ assay was 2.57 µg/mL (1.05 – 3.75 µg/mL) for a selectivity index of 10.54. These results showed that *U. tomentosa* known as Cat’s claw has antiviral effect against SARS-CoV-2 observed as a reduction in the viral titer and CPE after 48h of treatment on Vero E6 cells. Therefore, we hypothesized that *U. tomentosa* stem bark, could be promissory to the development of new therapeutic strategies against SARS-CoV-2.

## 1. Introduction

The severe acute respiratory syndrome coronavirus 2 (SARS-CoV-2) has caused serious public health problems since it was identified in Wuhan (China) in late 2019 [1]. The World Health Organization (WHO) declared Coronavirus disease 2019 (COVID-19) a pandemic on March 11, 2020 [2]. According to the latest report of the WHO, over 40,2 million cases and 1,100,000 deaths by COVID-19 were confirmed to October 20, 2020 [3]. When, the novel coronavirus (SARS-CoV-2) arrived to Latin-America, Brazil was the first South American country to declare a patient with COVID-19 whereas Venezuela and Uruguay were the ultimate nations to confirm their patient zero considering the pandemic epicenter after Europe [4]. Although some vaccines phase 3 medical trials are being tested in different countries sponsored by the pharmaceutical industry, currently, there is no vaccines, preventive treatment or antiviral drug available against SARS-CoV-2 [5].

Nowadays, the World Health Organization (WHO) recognizes that traditional, complementary, and alternative medicine has many benefits [6]. Several candidates with possible antiviral effects have been explored from medicinal plants in the preclinical phase. *Uncaria tomentosa* (Willd) DC., (*U. tomentosa*) belongs to Rubiaceae family, which is also known as Cat’s claw, it contains more than 50 phytochemicals [7]. Oxindole alkaloids (pentacyclic oxindole alkaloids (POA) and tetracyclic oxindole alkaloids (TOA)) have been recognized as fingerprint of this species in some pharmacopeias and several pharmacological activities are linked to this kind of alkaloids [8,9]. It has been demonstrated that *Uncaria tomentosa* (*U. tomentosa*) exerts an antiviral effect on human monocytes infected with dengue virus 2 (DENV-2) [10] and herpes simplex virus type 1 (HSV-1) [11]. In our previous studies *in silico, U. tomentosa’s* components inhibited the SARS-CoV-2 enzyme 3CL^pro^ and disrupted the interface of the receptor-binding domain of angiotensin-converting enzyme 2 (RBD–ACE-2) as well as the SARS-CoV-2 spike glycoprotein [12,13]. Additionally, bio-activities as anti-inflammatory [14], antiplatelet [15] and immunomodulatory [16] were demonstrated. Furthermore, other components isolated from the stem bark such as quinovic acids, polyphenols (flavonoids, proanthocyanidins, and tannins), triterpenes, glycosides and saponins were identified by instrumental methods [9,17–20].

The evaluation of natural compounds to inhibit SARS-CoV-2 in preclinical studies might lead to discover new antiviral drugs and for a better understanding of the viral life cycle. Several cell lines such as Human airway epithelial cells, Vero E6 cells, Caco-2 cells, Calu-3 cells, HEK293T cells, and Huh7 cells are considered the best models *in vitro* to determine the antiviral activity against SARS-CoV-2 [21]. Hence, the investigations of medicinal plants using specially Vero E6 cell to replicate SARS-CoV-2 and given that is highly expressed the ACE -2 receptor, in kidney tissue, some mechanism could be evaluated due to the characteristics of this culture medium.

Although the pathophysiology of COVID-19 is not completely understood, a severe inflammatory process has been associated with the severity and progression of the disease [22]. Therefore, the immune activation so far described during the course of the infection as well as the pulmonary injury could be ameliorated by *U. tomentosa* due to its traditional use as anti-inflammatory [23], in the folk medicine from South America during years.

Based on its antiviral activity on other ARN virus and our *in-silico* findings against SARS-CoV-2, we assayed the hydroalcoholic extract of *U. tomentosa* stem bark from Peru as potential antiviral agent *in vitro* against this novel coronavirus.

## 2. Material and method

### 2.1 Plant material

The *U. tomentosa* (Cat’s claw) used in this investigation is dispensed to patients of the Complementary Medicine Service of EsSalud (Social Health Insurance) in Peru for inflammatory disorders. The raw material (stem bark) of *U. tomentosa* was sourced from the Pharmacy Office of EsSalud in Ica, Peru. Next, the stem bark of *U. tomentosa* was transported to the Faculty of Medicine, of the Universidad Nacional Mayor de San Marcos (UNMSM, Lima, Peru) in order to obtain the hydroalcoholic extract.

### 2.2 Obtaining extract from plant material

One hundred grams of the raw plant material (stem bark) of *U. tomentosa* was powdered and extracted with 700 ml of 70% ethanol at room temperature for 7 days. Then, the extract was evaporated by using rotary evaporation to obtain a desiccated extract, which was stored at 4 °C until further use.

### 2.3 Preparation of stock solution of *U. tomentosa* extract

One mg of *U. tomentosa* hydroalcoholic extract was suspended in 1mL of DMSO. The solution was maintained at room temperature, protected from light until use. To prepare a working solution, the stock was diluted to 50mg/mL in DMEM supplemented with 2% Fetal Bovine Serum (FBS) (Final concentration DMSO 5%).

### 2.4 Cell lines and virus

Vero E6 epithelial cell line from *Cercopithecus aethiops* kidney was donated by Instituto Nacional de Salud (INS) (Bogotá, Colombia). Cells were maintained in Dulbecco’s modified Eagle medium (DMEM) supplemented with 2% FBS and 1% Penicillin-Streptomycin. Cultures were maintained at 37°C, with 5% CO_2_ Infections were done with a viral stock produced from a Colombian isolate of SARS-CoV-2 (hCoV-19/Colombia/ANT-UdeA-200325-01/2020).

### 2.5 Cell viability assays

The viability of Vero E6 in the presence of *U. tomentosa* extract was evaluated using a MTT (4,5-dimethylthiazol-2-yl)-2,5-diphenyl tetrazolium bromide) assay. Briefly, Vero E6 were seeded at cell density of 1.0×10^4^ cells/well in 96-well plates and incubated for 24h at 37°C in a humidified 5% CO_2_ atmosphere. After, 100 µL of serial dilutions (1:2) of *U. tomentosa* extract ranging from 3.1 to 50 µg/mL were added to each well and incubated for 48□h, at 37°C with 5% CO_2_. After incubation, supernatants were removed, cells were washed twice with Phosphate Buffered Saline (PBS) (Lonza, Rockland, ME, USA) and 30 µL of the MTT reagent (Sigma Aldrich) (2 mg/mL) were added. The plates were incubated for 2 hours at 37°C, with 5 % CO_2,_ protected from light. Then, formazan crystals were dissolved by adding 100 µL of pure DMSO to each well. Plates were read in a multiskan GO spectrophotometer (Thermo) at 570 nm. The average absorbance of cells without treatment was considered as 100% of viability. Based on this control the cell viability of each treated well was calculated. The treatment concentration with 50% cytotoxicity (The 50% cytotoxic concentration - CC_50_) was obtained by performing nonlinear regression followed by the construction of a concentration-response curve (GraphPad Prism). For MTT assay, 2 independent experiments with four replicates each experiment were performed (n=8).

### 2.6 Antiviral Assay

The antiviral activity of *U. tomentosa* extract against SARS-CoV-2 was evaluated with a pre-post strategy where the treatment was added before and after the infection. Briefly, Vero E6 cells were seeded at density of 1.0□× □10^4^ cells/well in 96-well plates and incubated for 24□h, at 37°C with 5% CO_2_. After incubation, 50uL of double dilutions of cat’s claw (3.1 – 25 µg/mL) were added to the cell monolayers during 1 h, 37 °C, 5 % CO_2_ Then, the treatment was removed, and cells were infected with SARS-CoV-2 stock at a multiplicity of infection (MOI) of 0.01 in 50uL of DMEM supplemented with 2% FBS. The inoculum was removed 1-hour post infection (h.p.i), replaced by 170 µL of cat’s claw dilutions and incubated for 48 h, at 37 °C with 5 % CO_2_. After, cell culture supernatants were harvested and stored at −80□°C for virus titration by plaque assay and TCID_50_ assay. The supernatant of infected cells without treatment was used as infection control. Chloroquine (CQ) at 50 µM was used as positive control for antiviral activity; 2 independent experiments with 3 replicates of each experiment were performed (n=6).

#### 2.6.1 Plaque assay for SARS-CoV-2 quantification

The capacity of *U. tomentosa* extract to decrease the PFU/mL of SARS-CoV-2 was evaluated by plaque assay on Vero E6 cells. Briefly, 1.0 x 10^5^ Vero E6 cells per well were seeded in 24-well plates for 24 h, at 37°C, with 5% CO_2_. Tenfold serial dilutions of the supernatants obtained from the antiviral assay (200uL per well) were added by duplicate on cell monolayers. After incubation during 1h, at 37°C, with 5% CO_2_, the viral inoculum was removed and 1 mL of semi-solid medium (1.5% carboxymethyl-cellulose in DMEM 1X with 2% FBS and 1% Penicillin-Streptomycin) was added to each well. Cells were incubated for 5 days at 37°C, with 5 % CO_2_. Then, cells were washed twice with PBS. After, cells were fixed and stained with 500 uL of 4 % Formaldehyde / 1 % Crystal violet solution for 30 minutes and washed with PBS. Plaques obtained from each condition were counted. The reduction in the viral titer after treatment with each concentration of *U. tomentosa* extract compared to the infection control is expressed as inhibition percentage. Two independent experiments with two replicates of each experiment were performed (n=4).

#### 2.6.2 TCID_50_ for SARS-CoV-2 quantification

The capacity of *U. tomentosa* extract to diminish the CPE caused by SARS-CoV-2 on Vero E6 was evaluated by TCID_50_ assay. Briefly, 1.2 x 10^4^ Vero E6 cells per well were seeded in 96-well plates for 24 h, at 37°C, with 5% CO_2_. Tenfold serial dilutions of the supernatants obtained from the antiviral assay (50 µL per well) were added by quadruplicate on cell monolayers. After 1h incubation, at 37°C with 5% CO_2_, the viral inoculum was removed and replaced by 170 µL of DMEM supplemented with 2% FBS. Cells were incubated for 5 days at 37°C, with 5 % CO_2_. Then, cells were washed twice with PBS, and then fixed and stained with 100 uL/well of 4 % Formaldehyde / 1 % Crystal violet solution for 30 minutes. Cell monolayers were washed with PBS. The number of wells positive for CPE were determined for each dilution (CPE is considered positive when more that 30% of cell monolayer if compromised). The viral titer of TCID_50_/mL was calculated based on Spearman-Käerber method. The reduction of viral titer after treatment with each concentration of *U. tomentosa* extract compared to infection control is expressed as inhibition percentage. A control of cells without infection and treatment was included. Two independent experiments with two replicates of each experiment were performed (n=4).

### 2.7 Statistical analysis

The median inhibitory concentration (IC_50_) values represent the concentration of the *U. tomentosa* extract that reduces virus particle production by 50%. The CC_50_ values, represent the cat’s claw solution concentration that causes 50 % cytotoxicity. The corresponding dose-response curves were fitted by non-linear regression analysis using a sigmoidal model. The calculated selectivity index (SI) represents the ratio of CC_50_ to IC_50_. All data was analyzed with GraphPad Prism (La Jolla, CA, USA) and data are presented as mean ± SEM. Statistical differences were evaluated via Student’s t-test or Mann–Whitney U test, a value of p□≤□0.05 was considered significant, with *p□≤□0.05, ** p□≤□0.01 and *** p□≤□0.001.

## 3. Results

### 3.1 The cell viability assay on Vero E6 cells in the presence of the *U. tomentosa* extract

The viability of Vero E6 cells in presence of *U. tomentosa* was higher than 90.0 % at concentrations of 25.0 µg/mL or lower, after of 48 h of incubation **(Figure 1**). Cell viability at 50.0 µg/mL was 17.3 %; for this reason, this concentration was not included in the antiviral assay. The CC_50_ calculated for *U. tomentosa* was of 27.1 µg/mL. Chloroquine at 50uM (positive control of inhibition) did not affect the viability of Vero E6 cells (**Figure 1**).

**Figure 1.**
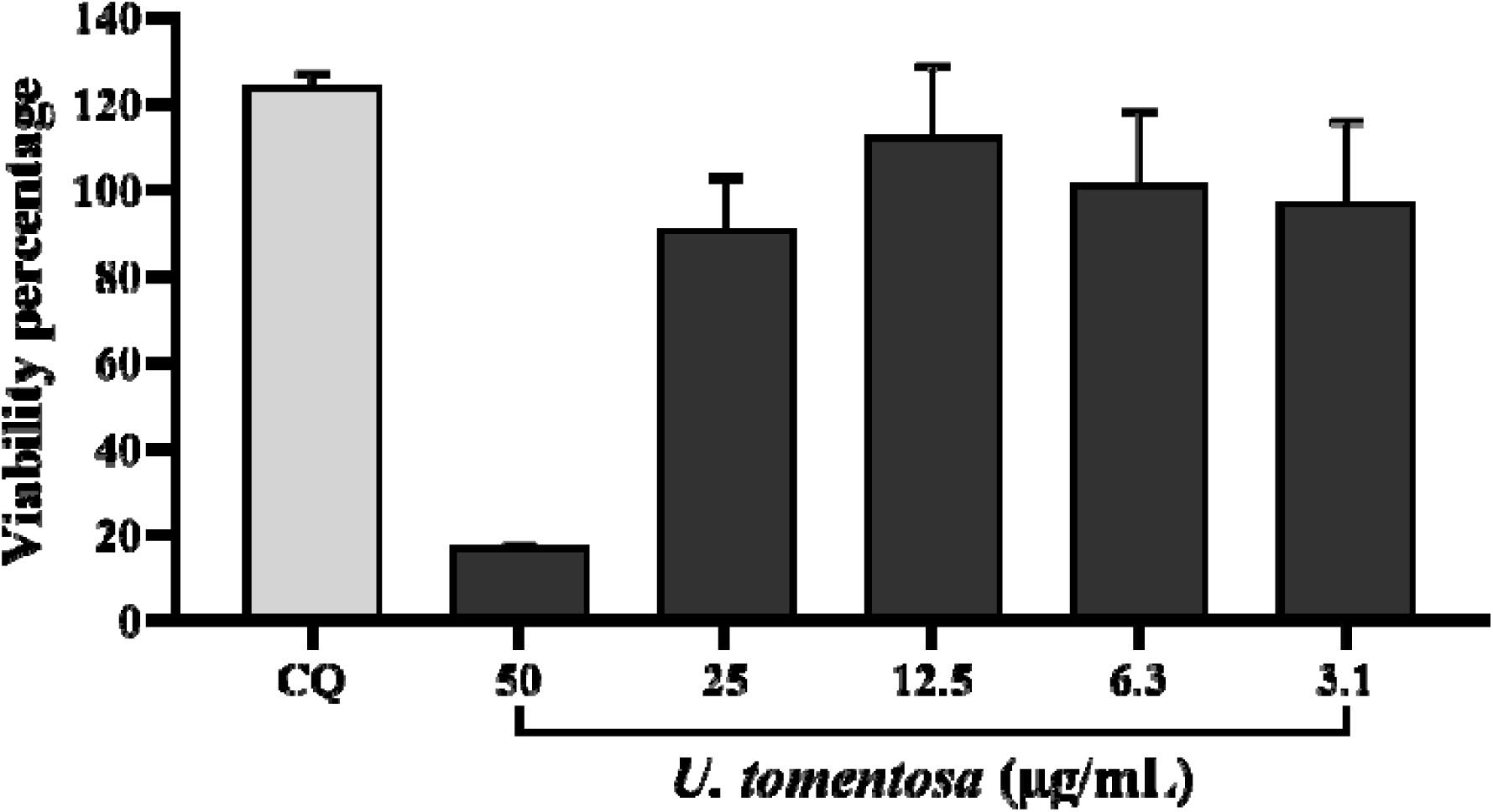
Vero E6 cells viability in presence of the *U. tomentosa* extract. The figure represents the viability percentage of Vero E6 cells after 48h of treatment with *U. tomentosa* (3.1 to 50.0 µg/mL). The viability percentages of treated cell were calculated based on the average absorbance control of cells without treatment. Chloroquine (CQ) was used as inhibition control of the antiviral strategy. Bars represent mean values ± SEM (2 independent experiments with four replicates each experiment were performed, n=8).

### 3.2 The *U. tomentosa* extract inhibited the number of infectious viral particles of SARS-CoV-2

An inhibition of 92.7 % of SARS-CoV-2 was obtained after the treatment with *U. tomentosa* at 25.0 µg/mL (p<0.0001) by plaque reduction assay (Figure 2). The *U. tomentosa* extract also showed an inhibition of 31.4 % and 34, 9 % of SARS-CoV-2 at 12.5 and 6.3 µg/mL, respectively (Figure 2). An increase of 76.0 % of PFU/mL of SARS-CoV-2 was obtained after the treatment with *U. tomentosa* extract at 3.1 µg/mL (p= 0.02) (**Figure 2**). The EC_50_ calculated to the extract by plaque assay was 6.6 µg/mL (4.89 – 8.85 µg/mL) for a selectivity index of 4.1. Chloroquine (inhibition positive control) showed an inhibition of 100 % of SARS-CoV-2 at 50 µM (p<0.0001) (Figure 2).

**Figure 2.**
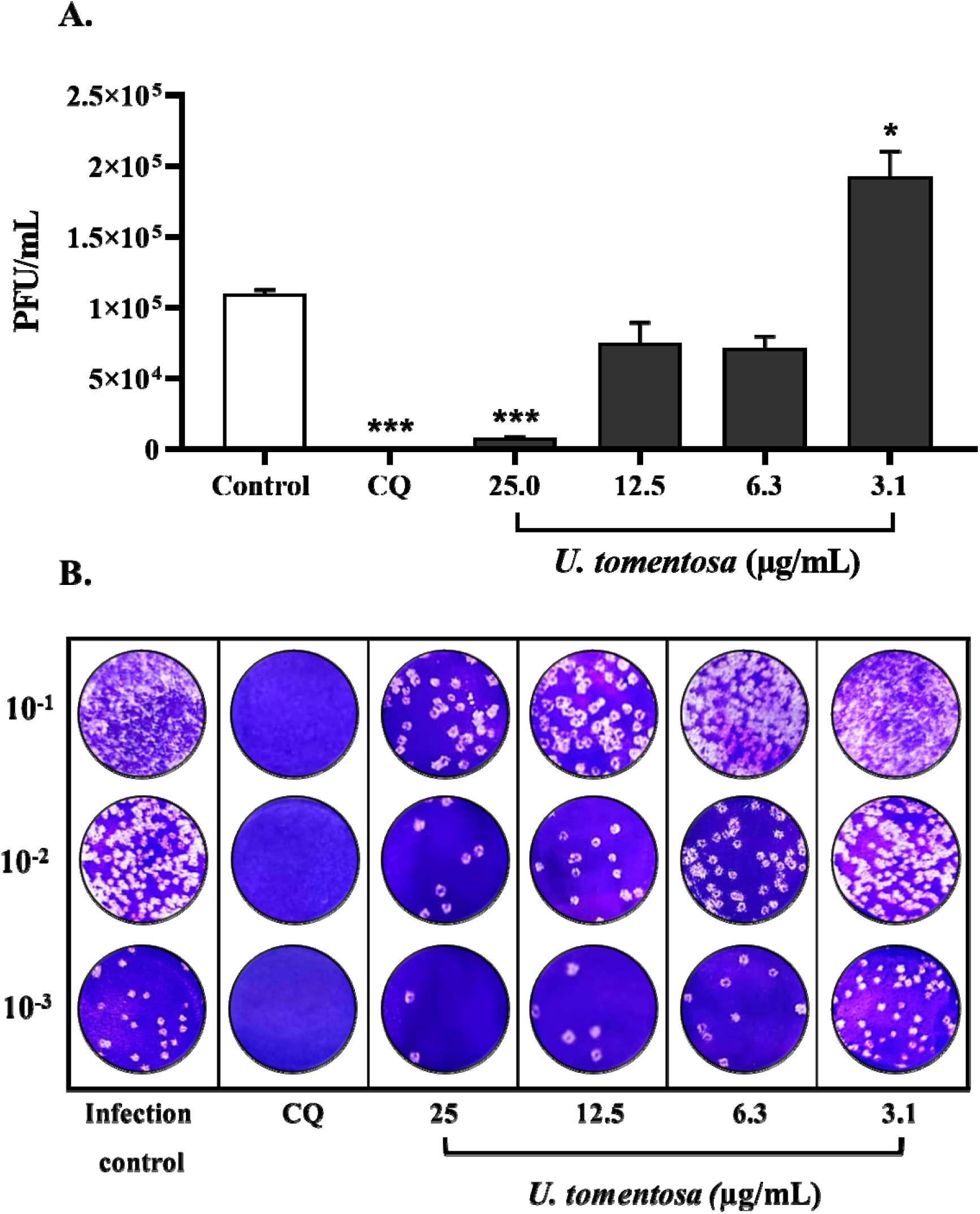
Antiviral activity *in vitro* of *U. tomentosa* extract against SARS-CoV-2 by plaque assay. **A**. The figure represents the viral titer (PFU/mL) of supernatants harvested after the treatment with the *U. tomentosa* extract quantified by plaque assay (n=4). Chloroquine (CQ) was used as an inhibition positive control of the antiviral strategy. * p□≤□0.05, ** p□≤□0.01 and *** p□≤□0.001 **B**. Representative plaques of the antiviral evaluation of the *U. tomentosa* extract against SARS-CoV-2 on Vero E6 cells.

### 3.3. The *U. tomentosa* extract reduced the CPE of SARS-CoV-2

The *U. tomentosa* extract reduced the CPE of the SARS-COV-2 on Vero E6 cells in a 98.6 % (p=0.02), 92.7 % (p=0.03), 63.2 and 60.4 % at 25, 12.5, 6.3 and 3.1 µg/mL, respectively (Figure 3). The EC_50_ calculated of *U. tomentosa* extract by TCID_50_ assay was 2.57 µg/mL (1.05 – 3.75 µg/mL) for a selectivity index of 10.54. Chloroquine (50 µM) inhibited the CPE of SARS-CoV-2 on Vero E6 in a 100 % (p=0.008) (**Figure 3**).

**Figure 3.**
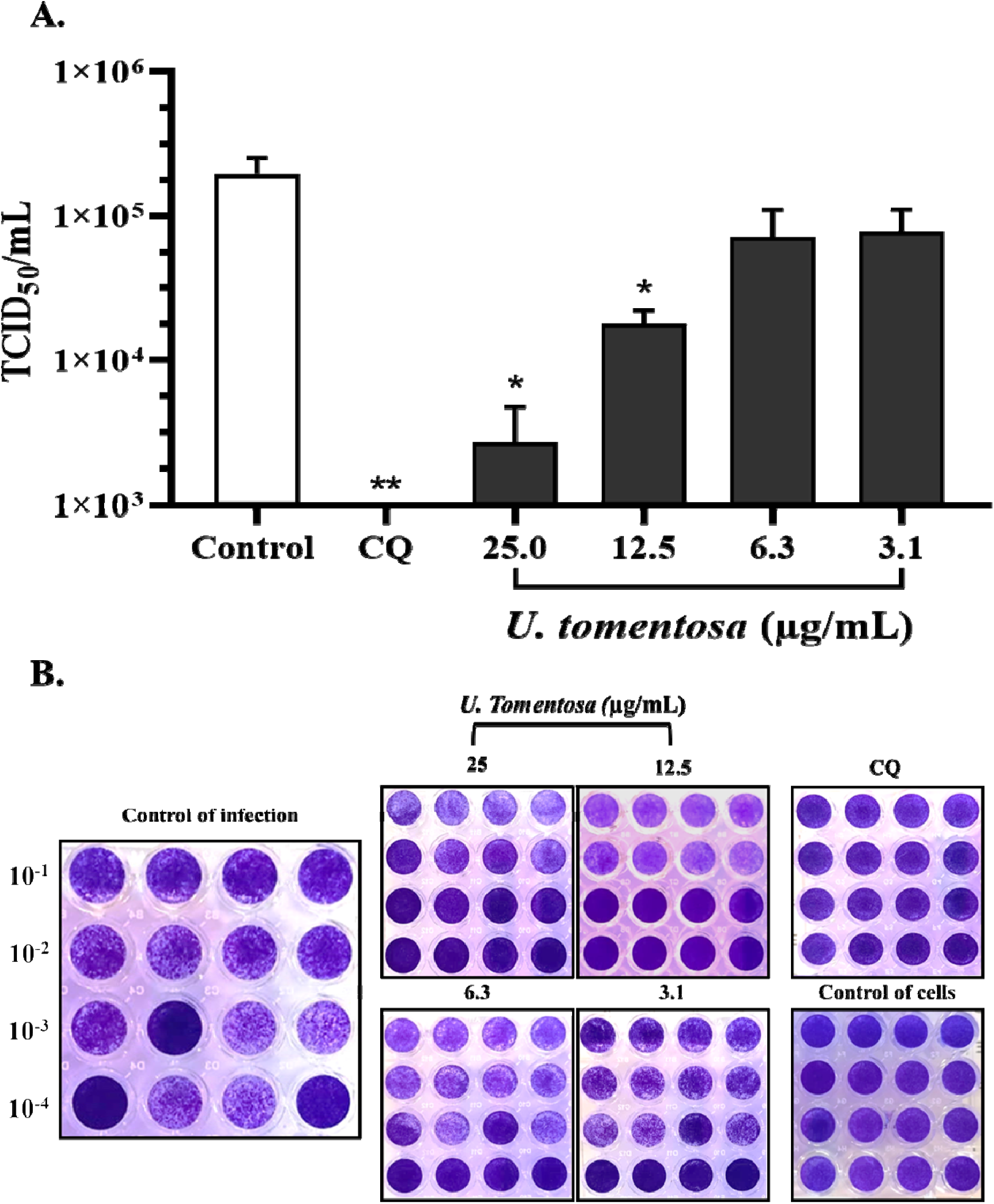
Antiviral activity *in vitro* of *U. tomentosa* extract against SARS-CoV-2 by TCID_50_ assay. The figure represents the viral titer (TCID_50_/mL) quantified by TCID_50_ assay on supernatants harvested from the treatment with the *U. tomentosa* extract (n=4). Chloroquine (CQ) was used as an inhibition positive control of the antiviral strategy. * p□≤□0.05 and ** p□≤□0.01 **B**. Representative images of the antiviral evaluation of the *U. tomentosa* extract against SARS-CoV-2 on Vero E6 cells by TCID_50_ assay revealed by crystal violet.

## 4. Discussion

In South America, the second wave of novel coronaviruses might be more aggressive, increasing the mortality rate and new cases [24]. Medical trials are underway to determine the efficacy of several vaccines against SARS-CoV-2 [25]. Otherwise, herbal medicines could become a promising option to tackle the ongoing pandemic caused by COVID□19 [26]. Some plant extracts and phytochemicals were modeled over numerous targets of SARS-CoV-2 by using *in silico* studies, which is the first step in the discovery of new drugs. In China, the use of herbal formulas has been included in the protocol of primary attention in COVID-19 and medical trials were carried out, and promising results to ameliorate the symptoms were demonstrated [27].

Our previous study of *U. tomentosa* (cat’s claw) on this novel coronavirus using *in-silico* analysis showed that two possible mechanisms could be involved in the *in vitro* antiviral activity against SARS-CoV-2 observed in this study. These findings revealed that 3CLpro, an essential enzyme for viral replication, showed key molecular interactions with speciophylline, cadambine, and proanthocyanidin B2, with high binding affinities ranging from −8.1 to −9.2□kcal/mol. [12]. On the other hand, phytochemicals of *U. tomentosa* such as Proanthocyanidin C1, QAG-2, Uncarine F, 3-isodihydrocadambine, and Uncaric acid (docking scores: -8.6; -8.2; -7.1; -7.6 and -7.0 kcal/mol, respectively) showed high binding affinity for the interface of the RBD–ACE-2. In addition, 3-dihydro-cadambine, Proanthocyanidin B4, Proanthocyanidin B2, and Proanthocyanidin C1 (−7.1; -7.2; -7.2 and -7.0 kcal/mol, respectively) had the highest binding score on SARS-CoV-2 spike glycoprotein [13]. Since Vero E6 cells are commonly used to replicate SARS-CoV-2 due to the high expression level of the ACE-2 receptor and lack the ability to produce interferon [28], they are the appropriate substrate to explore the antiviral activity of phytochemicals targeting the receptor binding.

Mechanisms of the antiviral activity of the hydroalcoholic extract of *U. tomentosa*, on other viruses like Dengue (DEN-2), have been elucidated; alkaloids (pentacyclic alkaloids) from *U. tomentosa* induced apoptosis of infected cells and reduced inflammatory mediators such as TNF-α, and IFN-α with similar effects to dexamethasone [10]. The quinovic acids (33.1-60 μg/mL) inhibited the Vesicular Stomatitis Virus [29], and the total extract at concentrations less than 15.75 μg/mL inhibited the Herpes Simplex Virus (HSV-1) replication when added to Vero cells at the same time than the virus [11].

Here, we demonstrated that *U. tomentosa* also has an antiviral activity *in vitro* against the SARS-CoV-2 with the EC_50_ calculated by plaque assay at 6.6 µg/mL (95% CI: 4.89 – 8.85 µg/mL) and by TCID_50_ assay at 2.57 µg/mL (95% CI: 1.05 – 3.75 µg/mL). Whilst the CC_50_ corresponding to a 50 % cytotoxic effect was 27.1 µg/mL for *U. tomentosa*. In other medicinal plants assayed against SARS-CoV-2, similar antiviral activity was shown; in particular, Echinaforce^®^ (an *Echinacea purpurea* preparation) exhibited an antiviral activity at 50 μg/mL [30]. Liu Shen capsule, a traditional Chinese medicine, inhibited the SARS-CoV-2 replication with an EC_50_ value of 0.6024 μg/mL and CC_50_ of 4.930 μg/mL [31]. Likewise, Phillyrin (KD-1), a representative constituent of *Forsythia suspensa* (Thunb.) presented an EC_50_ at 63.90 µg/mL and CC_50_ of 1959 µg/mL [32]. Sulfated polysaccharides named RPI-27 and heparin inhibited SARS-CoV-2 *in vitro* with an EC_50_ of 8.3 ± 4.6 μg/mL and 36 ± 14 μg/mL, respectively [33]. In our study, the selectivity index (SI) of the hydroalcoholic extract of *U. tomentosa* was 4.1. In spite of SI is a low value, theoretically having a higher value would be more effective and safer during *in vivo* treatment for a given viral infection. However, there is no evidence of severe toxicity of the *U. tomentosa* and traditionally its use popular in the form of maceration or decoction is safety [34].

There is enough evidence that *U. tomentosa* could ameliorate a wide array of symptoms associated with COVID-19, like the severe inflammation characterized by a cytokine storm [23] causing endothelial dysfunction. According to the antiviral activity of *U. tomentosa* against SARS-CoV-2, several biochemical mechanisms could be involved in each phase of the viral life cycle. As previously reported, *U. tomentosa* could interfere with viral entrance into host cells [12], affecting viral replication [13]. Furthermore, ACE-2 receptors, which are expressed in Vero E6 cells could also be blocked by the phytochemicals of *U. tomentosa* during the entrance of SARS-CoV-2 into the host cells and the aforementioned *in-silico* study would confirm our hypothesis [13].

Besides, it might control the hyperinflammation, via inhibition of IL-1α, 1β, 17, and TNF-α [35], reducing oxidative stress [36], and protecting the endothelial barrier, via inhibition of IL8, which is linked to the induction of permeability [37]. It also has antithrombotic potential via antiplatelet mechanism and by inhibiting of thrombin [15]. Furthermore, *U. tomentosa* modulates the immune system by extending lymphocyte survival via an anti-apoptotic mechanism [38]. It is known that the 3α protein of severe acute respiratory syndrome-associated coronavirus induces apoptosis in Vero E6 cells [39]; therefore, the phytochemicals found in the hydroalcoholic extract could inhibit this process and protect of the inflammatory cascade. Interestingly, *U. tomentosa* bark extract reduced the lung inflammation produced by ozone in mice [40].

Based on our results, *U. tomentosa* is a promising medicinal herb to combat COVID-19, but it is necessary to continue with animal models followed by clinical trials to validate our results in the context of COVID-19 patients. This study is the first approach of *U. tomentosa* against SARS-CoV-2 and we have to explore specific mechanisms of inhibition and propose the main molecules involved with the antiviral activity. As shown in our phytochemical analysis, the presence of groups chemicals determined by LC/MS (UHPLC-ESI+-HRMS-Orbitrap) such as spiroxindole alkaloids, indole glycosides alkaloids, quinovic acid glycosides, and proanthocyanidins, they could be responsible for the described activity. Here, the mechanisms discussed about the hydroalcoholic extract of *U. tomentosa* are only inferred under the mechanisms evaluated in other RNA viruses reported in the literature, and our previous *in-silico* studies demonstrated on the novel coronavirus.

## 5. Conclusion

*U. tomentosa* has been widely used as anti-inflammatory and immunomodulatory agent. Previous studies have shown that *U. tomentosa* has a broad spectrum of effects on several RNA viruses. In this study, we demonstrated that hydroalcoholic extract of *U. tomentosa* stem bark exerted an anti-SARS-CoV-2 activity by inhibiting virus replication *in-vitro* using the Vero E6 cell line, which might support to continue this investigation with specific assays *in-vitro*, then animal models and finally validate its clinical use with medical trials. However, further specific *in vitro* assays combined with *in vivo* studies need to be carried out to validate this *in-vitro* finding. Our investigation shows for the first time the antiviral effect of *U. tomentosa* on novel coronavirus (SARS-CoV-2).

## References

[1] Harcourt J, Tamin A, Lu X, Kamili S, Sakthivel SK, Murray J, et al. Severe acute respiratory syndrome coronavirus 2 from patient with coronavirus disease, United States. Emerg Infect Dis 2020. https://doi.org/10.3201/EID2606.200516.

[2] Takeda Y, Murata T, Jamsransuren D, Suganuma K, Kazami Y, Batkhuu J, et al. Saxifraga spinulosa-derived components rapidly inactivate multiple viruses including SARS-CoV-2. Viruses 2020. https://doi.org/10.3390/v12070699.

[3] World Health Organization. Coronavirus Dis Wkly Epidemiol Updat Wkly Oper Updat 2020. https://www.who.int/emergencies/diseases/novel-coronavirus-2019/situation-reports (Accessed October 4, 2020).

[4] Poterico JA, Mestanza O. Genetic variants and source of introduction of SARS-CoV-2 in South America. J Med Virol 2020. https://doi.org/10.1002/jmv.26001.

[5] Lai C-C, Shih T-P, Ko W-C, Tang H-J, Hsueh P-R. Severe acute respiratory syndrome coronavirus 2 (SARS-CoV-2) and coronavirus disease-2019 (COVID-19): The epidemic and the challenges. Int J Antimicrob Agents 2020;55:105924. https://doi.org/10.1016/j.ijantimicag.2020.105924.

[6] WHO. WHO supports scientifically-proven traditional medicineNo Title 2020. https://www.afro.who.int/news/who-supports-scientifically-proven-traditional-medicine (Accessed October 4, 2020).

[7] De Oliveira LZ, Farias ILG, Rigo ML, Glanzner WG, Gonçalves PBD, Cadoná FC, et al. Effect of Uncaria tomentosa extract on apoptosis triggered by oxaliplatin exposure on HT29 cells. Evidence-Based Complement Altern Med 2014. https://doi.org/10.1155/2014/274786.

[8] Heitzman ME, Neto CC, Winiarz E, Vaisberg AJ, Hammond GB. Ethnobotany, phytochemistry and pharmacology of Uncaria (Rubiaceae). Phytochemistry 2005. https://doi.org/10.1016/j.phytochem.2004.10.022.

[9] Lock O, Perez E, Villar M, Flores D, Rojas R. Bioactive compounds from plants used in peruvian traditional medicine. Nat Prod Commun 2016.

[10] Reis SRIN, Valente LMM, Sampaio AL, Siani AC, Gandini M, Azeredo EL, et al. Immunomodulating and antiviral activities of Uncaria tomentosa on human monocytes infected with Dengue Virus-2. Int Immunopharmacol 2008. https://doi.org/10.1016/j.intimp.2007.11.010.

[11] Caon T, Kaiser S, Feltrin C, de Carvalho A, Sincero TCM, Ortega GG, et al. Antimutagenic and antiherpetic activities of different preparations from Uncaria tomentosa (cat’s claw). Food Chem Toxicol 2014. https://doi.org/10.1016/j.fct.2014.01.013.

[12] Yepes-Pérez AF, Herrera-Calderon O, Sánchez-Aparicio J-E, Tiessler-Sala L, Maréchal J-D, Cardona-G W. Investigating Potential Inhibitory Effect of Uncaria tomentosa (Cat’s Claw) against the Main Protease 3CLpro of SARS-CoV-2 by Molecular Modeling. Evidence-Based Complement Altern Med 2020;2020:1–14. https://doi.org/10.1155/2020/4932572.

[13] Yepes-Pérez AF, Herrera-Calderon O, Quintero-Saumeth J. Uncaria tomentosa (cat’s claw): a promising herbal medicine against SARS-CoV-2/ACE-2 junction and SARS-CoV-2 spike protein based on molecular modeling. J Biomol Struct Dyn 2020:1–17. https://doi.org/10.1080/07391102.2020.1837676.

[14] Sandoval-Chacón M, Thompson JH, Zhang XJ, Liu X, Mannick EE, Sadowska-Krowicka H, et al. Antiinflammatory actions of cat’s claw: The role of NF-κB. Aliment Pharmacol Ther 1998. https://doi.org/10.1046/j.1365-2036.1998.00424.x.

[15] Kolodziejczyk-Czepas J, Ponczek M, Sady-Janczak M, Pilarski R, Bukowska B. Extracts from Uncaria tomentosa as antiplatelet agents and thrombin inhibitors – The in vitro and in silico study. J Ethnopharmacol 2020:113494. https://doi.org/10.1016/j.jep.2020.113494.

[16] Lenzi RM, Campestrini LH, Okumura LM, Bertol G, Kaiser S, Ortega GG, et al. Effects of aqueous fractions of Uncaria tomentosa (Willd.) D.C. on macrophage modulatory activities. Food Res Int 2013. https://doi.org/10.1016/j.foodres.2013.02.042.

[17] Navarro-Hoyos M, Lebrón-Aguilar R, Quintanilla-López JE, Cueva C, Hevia D, Quesada S, et al. Proanthocyanidin characterization and bioactivity of extracts from different parts of Uncaria tomentosa L. (cat’s claw). Antioxidants 2017. https://doi.org/10.3390/antiox6010012.

[18] Dietrich F, Kaiser S, Rockenbach L, Figueiró F, Bergamin LS, Cunha FM da, et al. Quinovic acid glycosides purified fraction from Uncaria tomentosa induces cell death by apoptosis in the T24 human bladder cancer cell line. Food Chem Toxicol 2014. https://doi.org/10.1016/j.fct.2014.02.037.

[19] Aquino R, De Tommasi N, De Simone F, Pizza C. Triterpenes and quinovic acid glycosides from Uncaria tomentosa. Phytochemistry 1997. https://doi.org/10.1016/S0031-9422(96)00716-9.

[20] Peñaloza EMC, Kaiser S, De Resende PE, Pittol V, Carvalho ÂR, Ortega GG. Chemical composition variability in the Uncaria tomentosa (cat’s claw) wild population. Quim Nova 2015. https://doi.org/10.5935/0100-4042.20150007.

[21] Takayama K. In Vitro and Animal Models for SARS-CoV-2 research. rends Pharmacol Sci 2020. https://doi.org/10.1016/j.tips.2020.05.005.

[22] Bohn MK, Hall A, Sepiashvili L, Jung B, Steele S, Adeli K. Pathophysiology of COVID-19: Mechanisms underlying disease severity and progression. Physiology 2020. https://doi.org/10.1152/physiol.00019.2020.

[23] Ferreira AO, Polonini HC, Dijkers ECF. Postulated Adjuvant Therapeutic Strategies for COVID-19. J Pers Med 2020;10:80. https://doi.org/10.3390/jpm10030080.

[24] Xu S, Li Y. Beware of the second wave of COVID-19. Lancet 2020. https://doi.org/10.1016/S0140-6736(20)30845-X.

[25] Rabaan AA, Al-Ahmed SH, Sah R, Tiwari R, Yatoo MI, Patel SK, et al. SARS-CoV-2/COVID-19 and advances in developing potential therapeutics and vaccines to counter this emerging pandemic. Ann Clin Microbiol Antimicrob 2020;19:40. https://doi.org/10.1186/s12941-020-00384-w.

[26] Ni L, Zhou L, Zhou M, Zhao J, Wang DW. Combination of western medicine and Chinese traditional patent medicine in treating a family case of COVID-19. Front Med 2020. https://doi.org/10.1007/s11684-020-0757-x.

[27] Xiong X, Wang P, Su K, Cho WC, Xing Y. Chinese herbal medicine for coronavirus disease 2019: A systematic review and meta-analysis. Pharmacol Res 2020. https://doi.org/10.1016/j.phrs.2020.105056.

[28] Ogando NS, Dalebout TJ, Zevenhoven-Dobbe JC, Limpens RWAL, van der Meer Y, Caly L, et al. SARS-coronavirus-2 replication in Vero E6 cells: Replication kinetics, rapid adaptation and cytopathology. J Gen Virol 2020. https://doi.org/10.1099/jgv.0.001453.

[29] Aquino R, De Simone F, Pizza C, Conti C, Stein ML. Plant metabolites. Structure and in vitro antiviral activity of quinovic acid glycosides from uncaria tomentosa and guettarda platypoda. J Nat Prod 1989. https://doi.org/10.1021/np50064a002.

[30] Signer J, Jonsdottir HR, Albrich WC, Strasser M, Züst R, Ryter S, et al. In vitro virucidal activity of Echinaforce®, an Echinacea purpurea preparation, against coronaviruses, including common cold coronavirus 229E and SARS-CoV-2. Virol J 2020. https://doi.org/10.1186/s12985-020-01401-2.

[31] Ma Q, Pan W, Li R, Liu B, Li C, Xie Y, et al. Liu Shen capsule shows antiviral and anti-inflammatory abilities against novel coronavirus SARS-CoV-2 via suppression of NF-κB signaling pathway. Pharmacol Res 2020. https://doi.org/10.1016/j.phrs.2020.104850.

[32] Ma Q, Li R, Pan W, Huang W, Liu B, Xie Y, et al. Phillyrin (KD-1) exerts anti-viral and anti-inflammatory activities against novel coronavirus (SARS-CoV-2) and human coronavirus 229E (HCoV-229E) by suppressing the nuclear factor kappa B (NF-κB) signaling pathway. Phytomedicine 2020. https://doi.org/10.1016/j.phymed.2020.153296.

[33] Kwon PS, Oh H, Kwon S-J, Jin W, Zhang F, Fraser K, et al. Sulfated polysaccharides effectively inhibit SARS-CoV-2 in vitro. Cell Discov 2020;6:50. https://doi.org/10.1038/s41421-020-00192-8.

[34] Maruoka T, Kitanaka A, Kubota Y, Yamaoka G, Kameda T, Imataki O, et al. Lemongrass essential oil and citral inhibit Src/Stat3 activity and suppress the proliferation/survival of small-cell lung cancer cells, alone or in combination with chemotherapeutic agents. Int J Oncol 2018. https://doi.org/10.3892/ijo.2018.4314.

[35] Rojas-Duran R, González-Aspajo G, Ruiz-Martel C, Bourdy G, Doroteo-Ortega VH, Alban-Castillo J, et al. Anti-inflammatory activity of Mitraphylline isolated from Uncaria tomentosa bark. J Ethnopharmacol 2012. https://doi.org/10.1016/j.jep.2012.07.015.

[36] Cheng AC, Jian CB, Huang YT, Lai CS, Hsu PC, Pan MH. Induction of apoptosis by Uncaria tomentosa through reactive oxygen species production, cytochrome c release, and caspases activation in human leukemia cells. Food Chem Toxicol 2007. https://doi.org/10.1016/j.fct.2007.05.016.

[37] Lima-Junior RS, Da Silva Mello C, Kubelka CF, Siani AC, Valente LMM. Uncaria tomentosa alkaloidal fraction reduces paracellular permeability, il-8 and ns1 production on human microvascular endothelial cells infected with dengue virus. Nat Prod Commun 2013. https://doi.org/10.1177/1934578X1300801112.

[38] Sandoval M, Charbonnet RM, Okuhama NN, Roberts J, Krenova Z, Trentacosti AM, et al. Cat’s claw inhibits TNFα production and scavenges free radicals: Role in cytoprotection. Free Radic Biol Med 2000. https://doi.org/10.1016/S0891-5849(00)00327-0.

[39] Law PTW, Wong CH, Au TCC, Chuck CP, Kong SK, Chan PKS, et al. The 3a protein of severe acute respiratory syndrome-associated coronavirus induces apoptosis in Vero E6 cells. J Gen Virol 2005. https://doi.org/10.1099/vir.0.80813-0.

[40] Cisneros FJ, Jayo M, Niedziela L. An Uncaria tomentosa (cat’s claw) extract protects mice against ozone-induced lung inflammation. J Ethnopharmacol 2005. https://doi.org/10.1016/j.jep.2004.06.039.

